# DNA Double Strand Break Repair in *E. coli* Perturbs Cell Division and Chromosome Dynamics

**DOI:** 10.1101/802389

**Authors:** M.A. White, E. Darmon, M.A. Lopez-Vernaza, D.R.F. Leach

## Abstract

To prevent the transmission of damaged genomic material between generations, cells require a system for accommodating DNA repair within their cell cycles. We have previously shown that *Escherichia coli* cells subject to a single, repairable site-specific DNA double-strand break (DSB) per DNA replication cycle reach a new average cell length, with a negligible effect on population growth rate. We show here that this new cell size distribution is caused by a DSB repair-dependent delay in completion of cell division. This delay occurs despite unperturbed cell size regulated initiation of both chromosomal DNA replication and cell division. Furthermore, despite DSB repair altering the profile of DNA replication across the genome, the time required to complete chromosomal duplication is invariant. The delay in completion of cell division is accompanied by a DSB repair-dependent delay in individualization of sister nucleoids. We suggest that DSB repair events create inter-sister connections that persist until those chromosomes are separated by a closing septum.

**Author Summary:** The bacterium *Escherichia coli* has a remarkable cell cycle where overlapping rounds of DNA replication can occur in a single generation between cell birth and division. This implies a complex coordination network between growth, genome duplication and cell division to ensure that the right number of genomes are created and distributed to daughter cells at all growth rates. This network must be robust to a number of unpredictable challenges. One such challenge is broken DNA, something that in *E. coli* is estimated to occur in ~20% of cell division cycles. In this work we perturb the *E. coli* cell cycle by elevating the frequency of repairable DNA double-strand breaks to determine which parameters of the cell cycle are conserved and which are changed. Our results demonstrate that this perturbation does not alter the average cell size at initiation of DNA replication or initiation of cell division. Furthermore, it does not alter the time taken to replicate the genome or the generation time. However, it does delay the segregation of the DNA to daughter cells and the completion of cell division explaining the increase in average cell size observed previously.

## Introduction

The presence of a 246bp interrupted DNA palindrome inserted at the *lacZ* locus of an otherwise wild-type *Escherichia coli* chromosome results in a chronic replication-dependent DNA double-strand break (DSB) that is efficiently repaired by homologous recombination with an unbroken sister chromosome [1]. This DSB is caused by the Mre11/Rad50 related endonuclease SbcCD cleaving a hairpin structure formed by the palindrome on one of a pair of sister chromosomes during DNA replication (Fig 1A). Cells undergoing this type of DNA double-strand break repair (DSBR) event are, on average, longer than control cells lacking either the palindrome or SbcCD [2]. This new cell length is accompanied by no detectable change in growth rate and a very small effect on cellular fitness (0.6% loss of fitness per generation) that is only detectable in co-cultivation experiments. Importantly, the observed increase in cell length is primarily due to an increase in unit cell size that is independent of the well-described DNA damage-induced inhibitor of cell division SfiA [2]. This DSB therefore acts as a defined perturbation of the cell cycle that maintains growth rate while altering cell size. It is also biologically relevant, with efficiently repaired replication-dependent DSBs estimated to occur in approximately 1 in every 5 normal *E. coli* cell division cycles [3].

**Figure 1.**
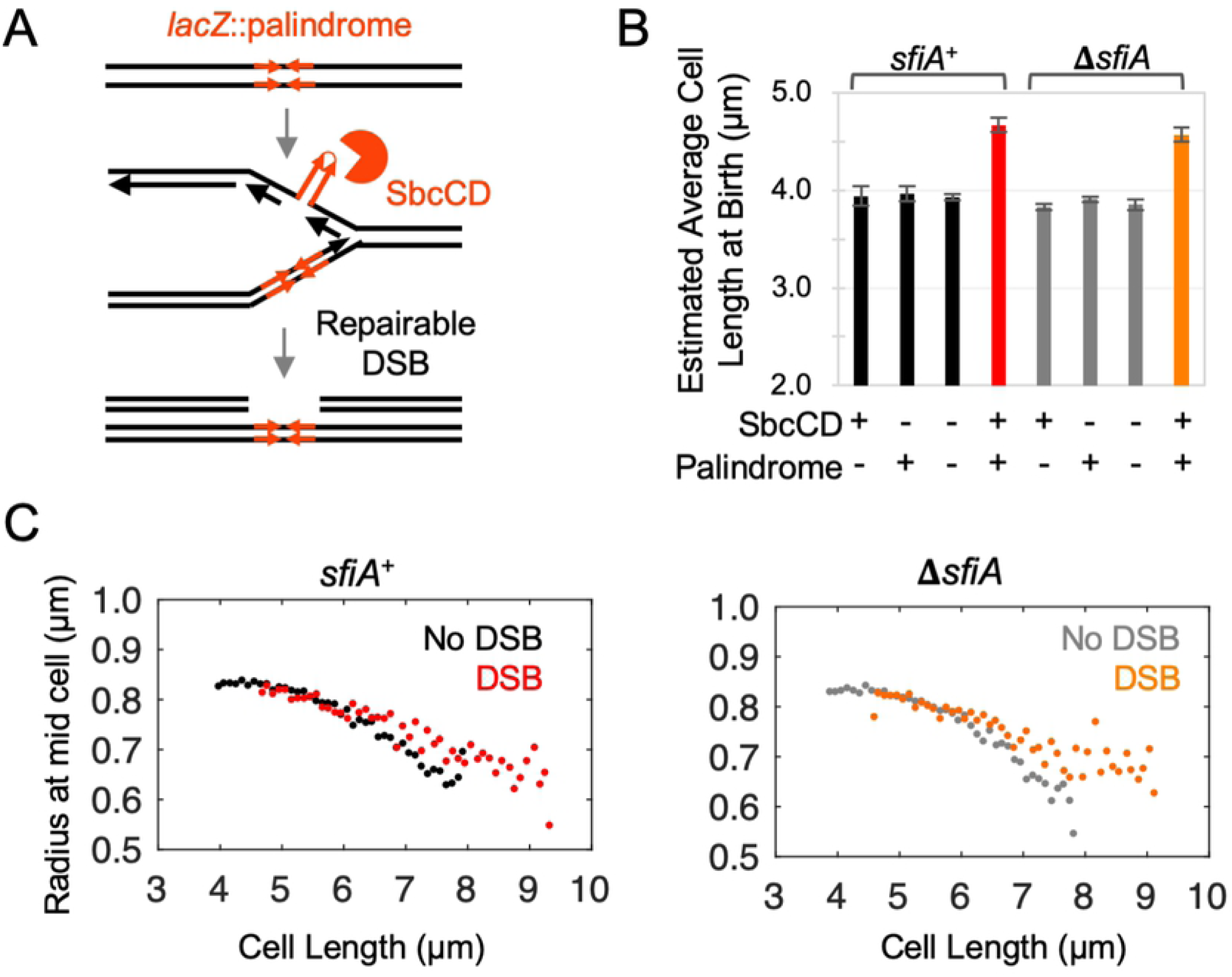
DSBR delays completion of cytokinesis without affecting the size control of its initiation. **A)** *E. coli* cells with an exogenous 246 bp interrupted DNA palindrome integrated into the *lacZ* gene of their chromosome undergo DNA double-strand break repair (DSBR) by homologous recombination once per replication cycle due to replication-dependent cleavage of the palindrome by the hairpin endonuclease SbcCD. **B)** Estimated average length at birth assuming an idealized population structure of cultures of cells undergoing DSBR (SbcCD^+^ *lacZ*::palindrome) and controls; n = 4, error bars show standard error of the mean. **C)** Radius of mid-cell (the location of cytokinesis) plotted as a function of cell length. Data from four independent cultures were aggregated and mid-cell radii averaged within 100 nm cell length. For both *sfiA*^+^ and Δ*sfiA* strains, the data from the three control strain backgrounds not undergoing DSBR (SbcCD^+^ *lacZ*^+^, SbcCD^-^ *lacZ*^+^ and SbcCD^-^ *lacZ*::palindrome) were averaged for clarity in the plots. No difference was detected between the three control strains in either *sfiA*^+^ or Δ*sfiA* genetic backgrounds.

*E. coli* has evolved a distinct cell cycle program where the generation (*i.e.* inter-division) time is not limited by the time required to replicate and segregate its genomes. This plasticity is achieved by the ability to initiate a new round of chromosomal DNA replication prior to completion of the previous round. This occurs in wild-type cells under fast growth conditions (mass doubling time less than ~ 60 min). It was initially observed by Cooper and Helmstetter, who termed the strategy “multi-fork replication” and proposed a simple model that provides a theoretical framework for understanding how *E. coli* can transition from a eukaryotic-like cell-cycle (one round of DNA replication within one cell division cycle) at slow growth rates to multi-fork replication at fast growth rates [4]. The underlying logic of this model is that replication initiates at regular intervals (defined by the generation time but independent of cell division) and that cell division occurs in a deterministic manner ~60 minutes following this event. Cooper and Helmstetter noted that this interval was longer than the time required for cells to complete chromosomal replication (the ‘C period’) and they named the remainder of this time the D period.

Based on a combination of measurements made in *E. coli* and *Salmonella typhimurium*, Donachie proposed that the underlying signal for periodic DNA replication initiation was the achievement of a critical cell mass [5], providing both a possible mechanism for coupling cell growth, genome duplication and cell division, as well as a plausible explanation for cell size homeostasis. Although doubt has been cast on the existence of a fixed critical cell mass for all growth conditions [6], the available data is consistent with cell-size regulated initiation of chromosomal DNA replication [7-9]. It is less clear however, whether or not (and if so, how) the events of replication initiation and cell division are causally linked [10-13].

By using a combination of whole genome sequencing, flow cytometry and imaging, we provide here a detailed analysis of the effect of a repairable DNA lesion on the cell cycle. We conclude that three key aspects of the DNA replication cycle (the average cell length at initiation of cytokinesis, the average cell length at initiation of DNA replication and the average duration of chromosomal replication) are maintained approximately invariant despite the perturbation of chronic DSBR. On the other hand, the individuation of nucleoids and the completion of cell division are delayed. We discuss two alternative, but not mutually exclusive, hypotheses for the impact of DSBR on the cell cycle. Either DSBR events in the current cell cycle are causing the delay in the current division event (inter-sister repair has an impact on inter-cousin segregation), or DSBR events create inter-sister chromosome connections that persist for several DNA replication cycles until those chromosomes are separated by a closing septum. The second hypothesis is more attractive, as it provides a plausible molecular explanation for both the impact of DSBR on nucleoid individuation and on cell division.

## Results

### DSBR delays completion of cytokinesis without affecting the size control of its initiation

Asynchronous cultures of *E. coli* cells were grown to exponential phase under conditions leading to an average mass doubling time of 17.4 min (Fig S1A, B), whereupon they were sampled for microscopy (Fig S1C). As previously reported, DSBR caused by the presence of a 246 bp interrupted DNA palindrome in cells expressing SbcCD (Fig 1A) resulted in a SfiA-independent increase in mean cell length (Fig S1D) without changing average mass doubling time (Fig S1B), [2]. Assuming an idealized population structure [6, 7, 14], Fig S1E-G), this translates to a SfiA-independent, DSBR-induced increase in average cell size at birth of approximately 18% (Fig 1B).

This DSBR-induced delay in cell division could be caused by either a delay in initiation of cell division, a delay in completion of cell division, or a combination of the two. *E. coli* are rod shaped cells that divide symmetrically to give two daughter cells. Progression of cytokinesis (septation) is therefore a function of cell radius at mid-cell [15]. *E. coli* cells exert cell size homeostasis and cell cycle control may be issued in response to either a time or size-dependent signal (see introduction).

Contrary to expectations for a perturbation that delays cell division, plotting cell radius at mid-cell as a function of estimated time since birth (Fig S1H) indicates that septation initiates relatively earlier in the cell division cycle in cells undergoing DSBR as the two curves diverge from cell birth. In contrast, plotting cell radius at mid-cell as a function of measured cell length (Fig 1C) indicates that DSBR affects cell division late in the septation process, once cells reach a length of ~6 µm. This indicates that septation initiation is either directly or indirectly initiated by a cell size-dependent mechanism (as opposed to, for example, time since cell birth Fig S1H) and that DSBR has no effect on septation initiation. Due to the structure of the population (there are twice as many new-born cells as cells about to divide), the data corresponding to the end of the cell division cycle are noisier than the data corresponding to the start of the cell division cycle (Fig 1C). As a result of this noise, it is not possible with the current dataset to distinguish whether the delay in completion of septation is due to a pause in septation and/or a decrease in the rate of septation.

### DSBR alters the chromosomal DNA replication profile without affecting the size control of replication initiation or the time required to complete DNA synthesis

Since DSBR is occurring on chromosomal DNA and chromosomal DNA is known to have the capacity to interfere with cytokinesis, we hypothesized that DSBR perturbs the chromosome cycle (replication initiation, completion and/or segregation).

### DSBR does not affect the cell size control of initiation of chromosomal DNA replication

Initiation of chromosomal DNA replication in *E. coli* is a cell size regulated process [5, 6, 9, 16]. To test if DSBR affects initiation of chromosomal DNA replication, cells were grown under the same conditions used above and the per-cell copy number of the origin of replication *oriC* was measured by ‘replication runout’ [6]. The observed DSBR-dependent increase in the relative proportion of cells in the asynchronous population that had initiated replication at the time of sampling (Fig 2A) indicated that DSBR caused chromosomal DNA replication to be initiated (on average) earlier in the cell division cycle. As observed for initiation of cytokinesis (above), this may or may not be due to the DSBR-dependent increase in average size at birth. The estimated average cell length at replication initiation (Fig 2B) can be calculated using both the estimated average size at birth (Fig 1B, above) and the measured relative fraction of the cell division cycle at which replication initiation occurs (age at initiation, A_i_, Fig 2A, [6]). This confirms that chromosomal DNA replication is initiated directly or indirectly by a size-dependent mechanism and indicates that DSBR does not affect this process (Fig 2B). For cells growing at the same average rate, initiating DNA replication at the same average size but dividing at a larger average size implies extending the time between initiation of DNA replication and completion of cell division (referred to as C+D time in the Cooper-Helmstetter model for the *E. coli* cell cycle), and this is indeed the case here where C+D is extended by 3.5 - 4 min in cells undergoing DSBR (Fig 2C). Assuming a baseline of the average division length of the control cells, the measured extension to C + D time predicts an average birth length of 4.63 µm and 4.57 µm for *sfiA*^+^ and Δ*sfiA* genotypes undergoing DSBR. This is in close agreement with the estimated average length of birth derived from the observed mean length and assuming an ideal population structure (4.67 µm and 4.52 µm for *sfiA*^+^ and Δ*sfiA* genotypes undergoing DSBR, see above, Figure 1B), demonstrating that the increased cell size caused by DSBR can, within the accuracy of the experiments, be accounted for by the increased time taken between initiation of replication and cell division.

**Figure 2.**
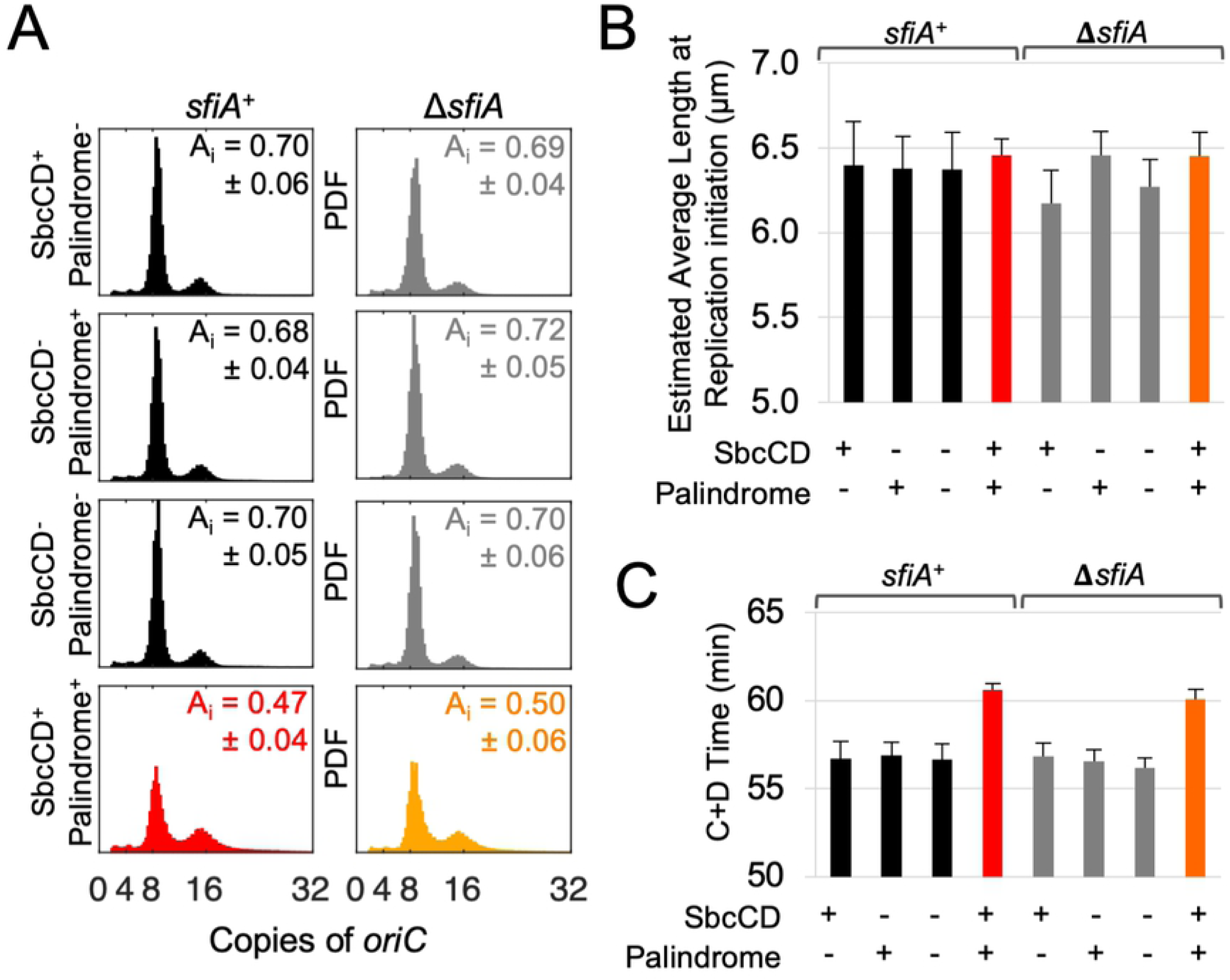
DSBR does not affect the cell size control of initiation of chromosomal DNA replication. **A)** Distribution of the copy number of the single chromosomal origin of replication (*oriC*) in cells isolated from asynchronous, exponentially growing cultures of *E. coli* as ascertained by replication runout. Shown are representative results from three independent experiments for each strain. Data are normalized as Probability Distribution Functions (PDFs). Insert of each panel shows the estimated relative cell age at replication initiation (A_i_), +/- the standard error of the mean. **B)** Estimated average cell length at initiation of chromosomal DNA replication calculated using the measured average copy number of *oriC* from three independent experiments and the estimated average cell length at birth (Fig 1B). **C**) C+D time (a component of the Cooper-Helmstetter model) calculated using the average mass doubling time for all strains (Fig S1B) and the average cellular *oriC* copy number for each strain (**A**). For B and C, error bars show standard error of the mean.

### DSBR changes the profile of chromosomal DNA replication

The observed delay in cell division relative to the initiation of DNA replication led us to investigate how DNA replication is perturbed by DSBR and in particular whether it takes longer to replicate genomes undergoing DSBR. Under the fast growth conditions used here, *E. coli* cells are continuously replicating their circular chromosomal DNA throughout the cell division cycle. The relative abundance of chromosomal loci across the genome in asynchronous cultures of *E. coli* cells is a function of the rate of replication and the generation time [17], with loci proximal to the origin of replication (*oriC*) being more abundant than loci close to the replication terminus. Marker frequency analysis (MFA) revealed this to be true for cultures of cells undergoing DSBR and controls (Fig 3A, S2A). DSBR-dependent changes in copy number across the genome (a combination of both genome-wide changes in DNA replication and local DNA processing that occurs at the site of the DSB) was revealed by normalizing the plots for cultures undergoing DSBR to controls (Fig 3B, S2B).

**Figure 3.**
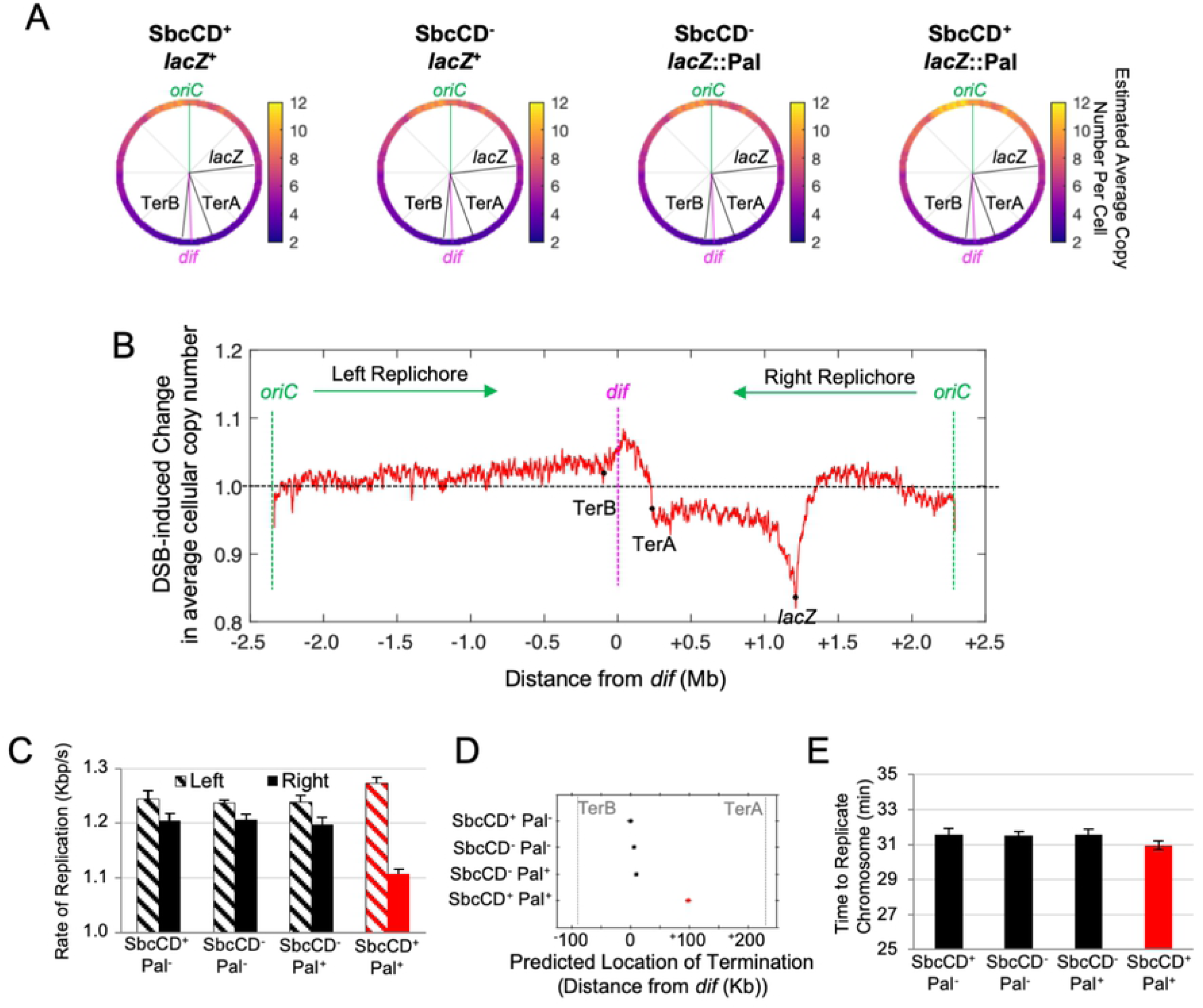
DSBR alters the chromosomal DNA replication profile without affecting the time required to complete DNA synthesis. **A)** Average copy number of chromosomal loci per cell across the 4.6 Mb circular genome in asynchronously growing cultures of *sfiA*^+^ cells. Average copy number per cell of each position in the genome was derived using the independently ascertained average copy number per cell of *oriC* (Fig 2A). Data shown are the mean from three independent experiments for each strain, each of which was normalized against a reference non-replicating (stationary phase) culture. **B)** Marker frequency analyses of cultures undergoing DSBR (SbcCD^+^ *lacZ*::palindrome) normalized against the average marker frequency of the three control strains not undergoing DSBR (SbcCD^+^ *lacZ*^+^, SbcCD^-^ *lacZ*^+^ and SbcCD^-^ *lacZ*::palindrome). **C)** Estimated average rate of replication of the two replication forks that initiate at *oriC*, calculated using the rate of change in marker frequency across the two halves (replichores) of the chromosome from three independent experiments. **D)** Estimated location of collision of the two replication forks that initiate at *oriC.* (**E**) Estimated average time to complete chromosomal DNA replication assuming DSBR-induced ectopic initiation of chromosomal replication in the Terminus region of the chromosome. For **C** - **E**, error bars show standard error of the mean, n = 3.

Cultures undergoing DSBR had, on average, more copies of sequences across the left half of the chromosome (‘left replichore’) than controls and this difference increased as a function of distance from *oriC* to the replication terminus. This is consistent with a DSBR-dependent increase in the rate of replication across the left replichore. The DSBR-induced changes in marker frequency across the right replichore are dominated by a local reduction of marker frequency flanking the locus undergoing DSBR (*lacZ*, Fig 3B) and a loss of reads that extends from the DSB locus all the way to the Terminus region. This is consistent with a RecBCD-mediated loss of reads in the vicinity of the DSB and a delay in the replication of the origin-distal arm of the chromosome from the DSB to the terminus.

The profile then changes in the Terminus region (Fig 3B). The ratio of reads in cells undergoing DSBR to control cells peaks between TerB and TerA (unidirectional terminator sequences that bind the replication fork blocking protein Tus). This is indicative of DSBR-induced ectopic initiation of DNA replication in the terminus. It is currently unclear what the source of this DSBR-dependent increase in terminus copy number is, although initiation of chromosome replication within the Terminus region has been previously reported [18].

### The time required to replicate chromosomal DNA is similar despite DSBR

The population average rate of DNA synthesis for both replication forks initiated from *oriC* can be derived from the rate of change of copy number from *oriC* to the replication terminus ([17] Fig S2C). In the absence of DSBR, each of the two replication forks were found to replicate chromosomal DNA at a rate of ~1.2 kb/s, with the left arm of the chromosome being replicated on average at a slightly faster rate than the right arm of the chromosome (Fig 3C). In cultures undergoing DSBR at *lacZ*, the rate of replication across the right arm of the chromosome was reduced by ~10% to 1.1 kb/s. A concomitant increase in the rate of replication across the left replichore was also observed (Fig 3C).

The population average location of replication termination can be derived by determining the point of interception of the lines of best fit to both replichores (Fig S2D). Consistent with published data on GC / AT sequence bias [19], replication was found to terminate, on average, close to the *dif* locus in the absence of DSBR (Fig 3D). This method assumes that replication initiates at a single locus (*oriC*). Determination of the average location of replication termination in cultures of cells undergoing DSBR is complicated by what appears to be DSBR-dependent ectopic initiation of DNA replication in the terminus region and by the MFA data representing an average behavior of cells across the whole population. It is possible that there are two populations of cells in cultures undergoing DSBR. In one population, there is no initiation of replication in the terminus and the forks meet at the average position, 99kb from *dif* on the right replichore, determined by the replication rates of the two replichores (Fig 3D). In the second population, replication initiated in the terminus causes termination at TerA and/or TerB so reducing the origin-initiated replication by up to 267.4 kb (the distance between TerA and TerB).

The population average total time to complete DNA synthesis was derived using the rate of replication and the location of termination. In the absence of DSBR, chromosomal DNA replication was found to take 30 minutes to complete. Depending upon which of the two termination positions above was used, DSBR was determined to either reduce, or increase, the time required to complete DNA synthesis by 30 seconds. Figure 3E depicts the result obtained assuming ectopic initiation of chromosomal replication in the terminus. Neither difference was found to be statistically significant (p = 0.37 and p = 0.15 respectively, one-way analysis of variance). We therefore conclude that DSBR has little if any effect on the observed 3.5 - 4 min increase in C+D time, leading us to conclude that, in the context of the Cooper-Helmstetter model, DSBR primarily extends D time.

### DSBR delays segregation of chromosomal DNA from the division plane

Cells that were sampled for microscopy were treated with chloramphenicol and DAPl to visualize their chromosomal DNA. Chloramphenicol causes chromosomes to condense and is therefore likely to induce the individualization of nucleoid bodies lacking (stable) inter-nucleoid linkages. The DAPl signal from each cell was projected along its long axis to give a 1D readout of DNA intensity. Cells were sampled taking into account the population structure, ordered according to cell length and the signal aligned by mid-cell to produce a pseudo-time series representing the average cell division cycle (Fig 4).

**Figure 4.**
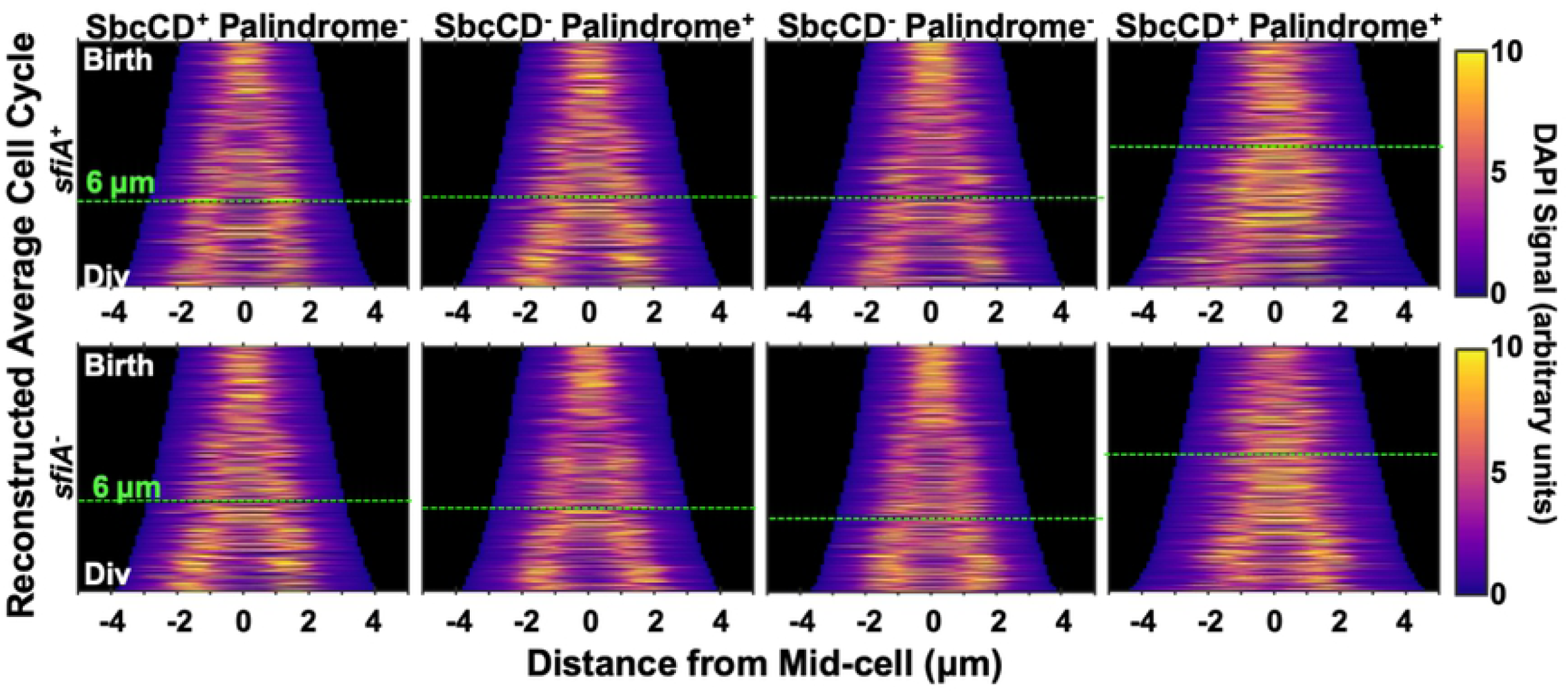
Cells undergoing DSBR have unsegregated chromosomal DNA at the division plane at the time of the block to cytokinesis. Intensity of DAPl signal from cells treated with DAPl and chloramphenicol projected along the long axis of cells and plotted as a function of cell length. For each strain, only cells between the estimated length at birth and division (Fig 1B, S1G) were included. Average cell division cycles were reconstructed by taking the age structure of the population (Fig S1F) into account when sampling of cells.

In the absence of induced DSBR, sister nucleoids appear to segregate and merge several times during the cell cycle before segregating one final time prior to cell division. It is also possible that each individual cell has a single transition event from 1 to 2 nucleoids and that this transition event varies greatly between cells. In cultures of cells undergoing DSBR, both the time and size of the first detectable nucleoid individualization event are increased relative to the controls (Fig 4). This DSBR-correlated delay in nucleoid individualization occurs prior to, and extends beyond, the observed block to cytokinesis (at ~6µm cell length, Fig 1C).

### DSBR is concurrent with the block to cytokinesis, but does not cause the chromosomal locus undergoing repair to dwell at the division plane

DSBR by homologous recombination involves the physical interaction between two DNA molecules (the broken chromosome and the intact chromosome used as a template for repair). The average cell length at the time of *lacZ* replication (when the DSB forms) was estimated using the average mass doubling rate (Fig S1B), the estimated average cell length at replication initiation (Fig 2B) and the rate of DNA synthesis (Fig 3C) and found to be coincidental with the observed block to cytokinesis (~6 µm, Fig 1C, Fig 5A).

**Figure 5.**
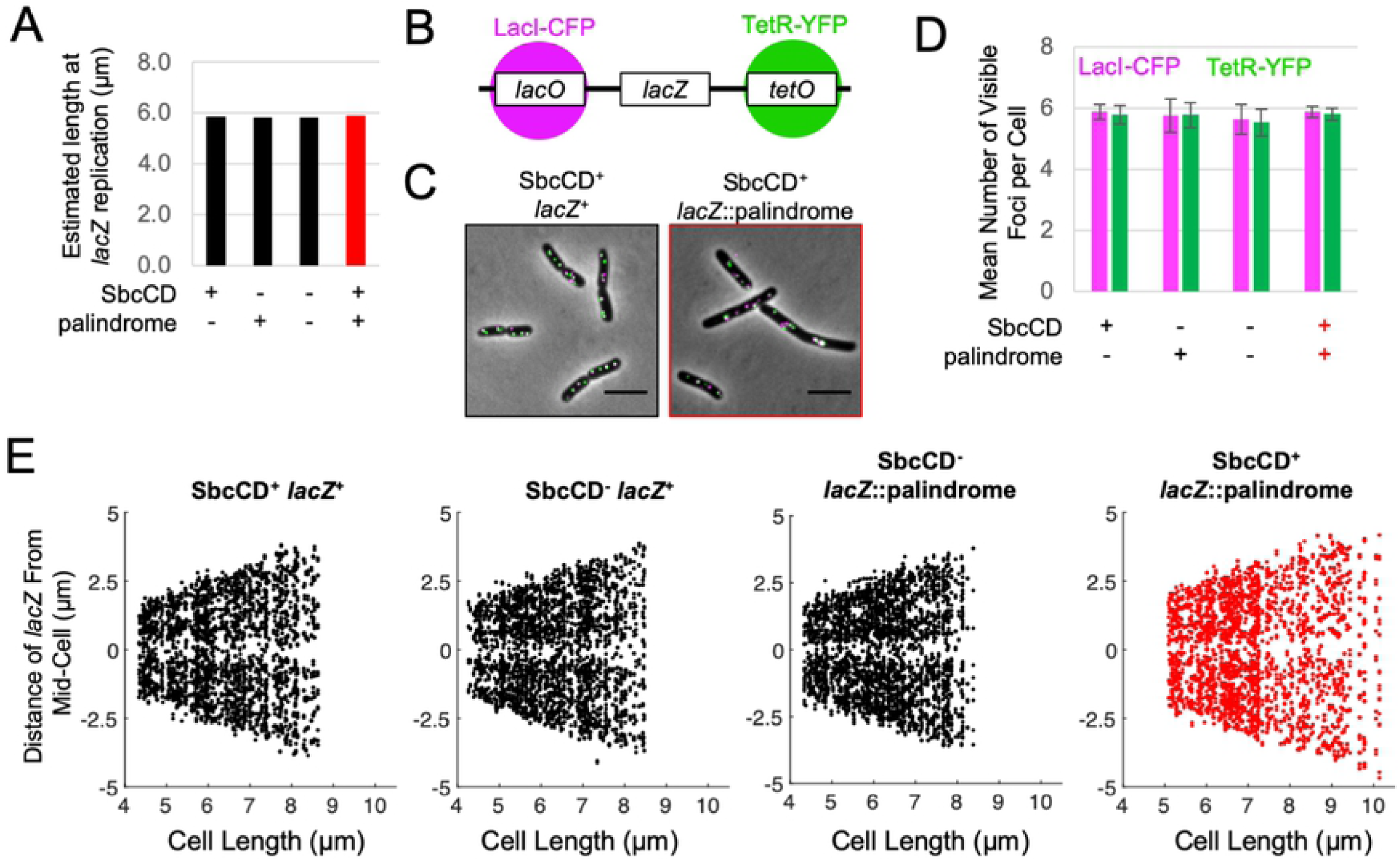
The locus undergoing DSBR does not dwell at the division plane. **A**) The average cellular length at the time of *lacZ* replication as estimated using the average mass doubling rate (Fig S1B), the estimated cellular length at replication initiation (Fig 2B), and the rate of DNA synthesis (Fig 3C). **B)** Diagram of the chromosomal construct for visualizing the cellular location of *lacZ*. The arrays of *lacO* and *tetO* operator sites were located respectively ~6 kb on the origin-proximal and ~5 kb on the origin-distal sides of the palindrome in *lacZ*. Lacl-CFP and TetR-YFP were expressed from a synthetic constitutive promoter integrated in the chromosome at the *ykgC* locus. **C**) Example images of *lacZ* localization in cells undergoing DSBR at *lacZ* (SbcCD^+^ *lacZ*::palindrome) or not (SbcCD^+^ *lacZ*^+^). Scale bar shows 5 µm. **D**) Mean number of *lacZ* associated Lacl-CFP and TetR-YFP foci per cell. Error bars show standard error of the mean for four independent cultures. **E**) The spatial distribution of *lacZ* associated foci along the long axis of cells as a function of cell length. Each panel shows the position of *lacZ*-associated foci for 300 cells whose lengths lie between the estimated average length at birth and estimated average length at division for the indicated strain. For each cell, the data for Lacl-CFP and TetR-YFP were aggregated.

To test if the observed delay in chromosome segregation from the division plane was a result of DSBR occurring at the division plane, we used a strain where the DSB locus (*lacZ*) was tagged at a distance of 5 - 6kb on either side with an array of *tetO* and an array of *lacO* sequences whose cellular location can be detected by fluorescence microscopy upon expression of TetR and Lacl fused to fluorescent proteins (TetR-YFP and Lacl-CFP, Fig 5B, C) [20, 21]. On average, cells had 6 segregated *lacZ* loci (Fig 5D) irrespective of whether or not they were undergoing DSBR. This implies that under these growth conditions, the majority of cells are cycling between 4 and 8 copies of *lacZ*, consistent with the calculated average cellular *lacZ* copy number (Fig 3A). Since cells undergoing DSBR show an increase in average birth size (Fig 1B) but the same average size at initiation of DNA replication (Fig 2B) and the time to replicate the region between *oriC* and *lacZ* is similar (Fig 3), we predict that replication of *lacZ* occurs earlier in the cycle of cells undergoing DSBR. Therefore, the observation of no change in the average number of segregated *lacZ* foci implies that homologous loci co-localize for an extended period of the cell division cycle in cells undergoing DSBR as shown to also be the case under slower growth conditions [22]. This was associated with a slight increase in the probability of *lacZ* foci being located at ¼ and ¾ positions of the cell length, in cells undergoing DSBR compared to controls (Fig S3). These positions correspond to the expected locations of future cell division. The *lacZ* loci that have undergone DSBR are well separated from each other and from the division plane suggesting that *lacZ* recombination is not directly interfering with cell division (Fig 5E). This argues that an uncharacterized event downstream of *lacZ* recombination is interfering with nucleoid individuation and cell division.

## Discussion

### Average cell length at initiation of cytokinesis, average cell size at initiation of DNA replication and average duration to replicate the chromosome remain approximately invariant despite DSBR

The resilience of living organisms depends on their capacity to maintain approximately invariant properties despite perturbations in their internal and external environment. In this study we have investigated the cell cycle consequences of perturbing the bacterium *E. coli* with a controlled, slightly elevated level of repairable replication-dependent DNA double-strand breaks. This perturbation has previously been shown to increase average cell size without measurably affecting growth rate [2]. Here we show that, in addition to the approximate invariance in growth rate, average cell length at initiation of cytokinesis, average cell size at initiation of replication and average length of time to replicate the chromosome remain approximately invariant despite DSBR.

The fact that average cell length at both initiation of chromosomal DNA replication and cytokinesis are invariant despite a DSBR-dependent increase in average length at birth implies that both of these crucial cell cycle events are either directly or indirectly regulated by one or more size-sensing mechanisms. Furthermore, the size control of these events is not perturbed by DSBR. By not affecting homeostatic control of cell size regulation, chromosomal and cell division events remain coupled and a DSBR-dependent constitutive delay in cell division can be tolerated without affecting population growth rate. Since DSBR was found to affect neither size control of initiation of chromosomal DNA replication nor cell division, these results cannot distinguish whether or not these cell cycle events are linked by the same size-regulatory pathway. It is, however, conceivable that a causal relationship could be tested by investigating this perturbation using single cell measurements that exploit the cell-to-cell variability of these events.

It is likely that the average cell size at initiation of replication and at initiation of cytokinesis are triggered independently of the events associated with DSBR. The time required to replicate the genome is also invariant despite there being a local impact of DSBR on the pattern of duplication of the genome as revealed by MFA (Fig 3). Here we can see that there is a loss of sequencing reads in the vicinity of the DSBR event that are likely to reflect a combination of processing of the DSB by RecBCD and loss of sequencing reads caused by as yet unresolved Holliday junctions [23]. There is also a loss of sequencing reads between the DSBR site and the chromosome terminus that can most easily be explained by a delay in replication forks reaching this region. This may be due to approximately 50% of two-ended DSBs being converted to one ended DSBs by RecBCD promoted degradation of the DNA end between the DSB site and the replication fork that had passed the site of the palindrome responsible for chromosome cleavage [24]. Following such degradation, replication of this region of the chromosome has to wait for recombination of the origin proximal DNA end and re-establishment of a replication fork. Despite these local perturbations to DNA replication, the combination of a slightly elevated rate of replication on the unaffected replichore and/or terminus-specific replication can compensate for the impact of DSBR and the overall time to replicate the genome is approximately conserved. Evidence exists that the rate of DNA replication is determined by the intracellular pools of deoxyribonucleotide triphosphates (dNTPs) [25-27] and this may indeed be the primary mediator of homeostatic regulation of the time taken to replicate the genome. lf effective replication is compromised on the right replichore, this could result in an elevation of dNTP pools that would cause accelerated replication on the left replichore via a complex regulatory network [28]. The underlying mechanism of the terminus-specific DNA synthesis is yet unknown but there is a long history of detection of replication in the terminus region of the *E. coli* chromosome [18] that can become very prominent in certain mutant strains (e.g. in the absence of RecG [29-31]).

### DSBR delays the completion of cytokinesis and the segregation of ‘cousin’ chromosomes from the division plane

We have shown here that the time between the initiation of DNA replication and the cell division associated with that initiation event is extended for cells undergoing DSBR (Fig 2C). Since this is not caused by an extension of the time to replicate the chromosome (Fig 3E), it is the time between the completion of DNA replication and the cell division event splitting that set of two chromosome termini into two cells that is extended. This is not associated with a delay in the initiation of cytokinesis but rather with a delay in the completion of cytokinesis (Fig 1C), which is consistent with a chromosome segregation problem. Further evidence of a perturbation to chromosome segregation comes from the DSBR induced delay in the individuation of chloramphenicol condensed nucleoids (Fig 4).

Our flow cytometry, microscopy and replication profiles argue that, under the rapid growth conditions used here, cells are born with eight origins of replication, four *lacZ* loci and two termini; they divide with sixteen origins, eight *lacZ* loci and four termini. Each cell therefore divides with four completed chromosomes (defined by the number of copies of the terminus) and the two daughter cells each receives a pair of sister chromosomes (Fig 6A). The two sister chromosomes in one cell are cousins of the two sister chromosomes in the other cell so the division event separates cousin chromosomes. DSBR does not change this. What does change is the interconnection of these sequences, as revealed when the nucleoids are condensed with chloramphenicol. The cells undergoing DSBR have trouble separating their nucleoids and this is accompanied by a delay in the completion of cytokinesis. The number of *lacZ* foci are little affected by DSBR despite DNA replication initiating earlier in the cells undergoing DSBR. This implies that DSBR increases the cohesion of the recombining chromosomes, as shown previously for slow growing cells [22]. However, the *lacZ* foci that have undergone DSBR are well separated by the time the two chromosomes they were located in are separated by cell division. In fact, by the time they are separated by division, they have undergone two further rounds of replication and DSBR to generate eight loci and it is clear that these loci do not dwell in the division plane. So, these eight *lacZ* loci are located in cousin chromosomes that need to split into two sister nucleoids before cytokinesis can be completed, and it is this separation of sister nucleoids that is delayed by DSBR. See Fig 6A for the predicted chromosomal structures expected to be present at cytokinesis and Fig 6B for an overall depiction of the cell cycle as perturbed by DSBR.

**Figure 6.**
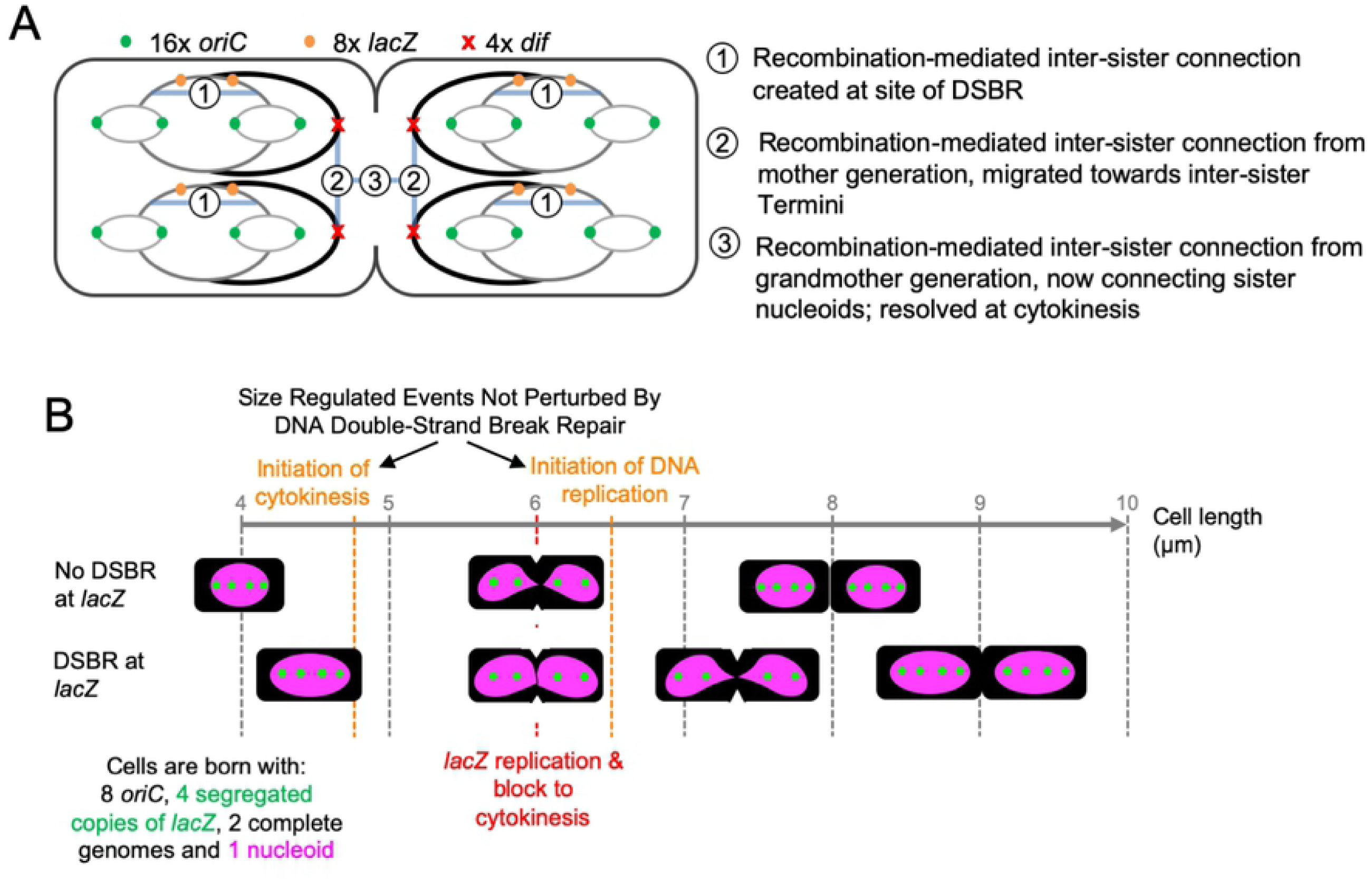
Diagrammatic representation of the cell cycle of fast-growing *E. coli* cells and its perturbation by DSBR at *lacZ*. **A)** Segregation of chromosomes in fast growing cells. At cytokinesis, there are four fully replicated chromosomal structures defined by the presence of four chromosome termini (*dif*). Each of these chromosomal structures contains partially re-replicated sections due to initiation of replication from the origin (*oriC*). Upon division, both halves of the cell will become a new-born cell, each containing two fully replicated sister chromosomal structures. At the time of cytokinesis, each of the four chromosomal structures contains a re-replicated *lacZ* locus and, in cells where DSBR has occurred, we hypothesize that a connection is generated between the two replicated and recombined double-stranded arms of DNA (shown by a blue band with the number one). We propose that the previous round of replication of *lacZ*, and its associated DSBR event, generated similar connections (depicted by blue bands with the number two) that interlink the sister chromosomal structures within the two halves of the cell. Similarly, when *lacZ* was replicated two rounds previously, connections (depicted by blue bands with the number three) were made between the cousin chromosomal structures attempting to segregate at this cytokinesis event. It is this latter set of chromosomal connections that we propose needs to be resolved in order for this cytokinesis event to be completed. The nature and number of the connections is unknown and so the blue bands do not represent specific numbers or locations of connections between the chromosomes. **B)** Implications of DSBR for cytokinesis. Fast-growing *E. coli* cells undergoing a normal cycle are born at a length of approximately 4 µm and divide at approximately 8 µm. They initiate cytokinesis at just under 5 µm, replicate the *lacZ* locus at approximately 6 µm and initiate DNA replication at *oriC* at about 6.5 µm. They are born with 8 copies of *oriC*, 4 segregated copies of *lacZ*, 2 complete genomes and 1 nucleoid, and divide with twice these constituents. Cells undergoing DSBR at *lacZ* are also born and divide with these numbers of copies of *oriC, lacZ*, genomes and nucleoids. Furthermore, they initiate cytokinesis, *lacZ* replication and *oriC* replication at the same cell sizes as the unperturbed cells. However, completion of cytokinesis is delayed until the cells are approximately 9 µm long and so they are born with a length of approximately 4.5 µm. Accompanying this delay to cell division is a delay to the individuation of nucleoids.

We have considered two alternative hypotheses to explain this behavior. Either DSBR events in one cell cycle cause a division delay in the same cell cycle, implying that inter-sister repair at the *lacZ* locus has an impact on inter-cousin chromosome segregation; or DSBR events at the *lacZ* locus create inter-sister chromosome connections that do not inhibit the separation of sister *lacZ* loci (following an initial delay) but persist for several DNA replication cycles and interlink cousin chromosomes until they are separated by a closing septum. The transmission of information from interacting sister *lacZ* loci to segregating cousin Terminus loci (the last locus to segregate during the *E.coli* cell cycle [32]) could be accomplished by physical forces that can propagate along the mechanically coherent nucleoid structure in the order of seconds [33]. Notably, due to the absence of an active mechanism for bulk nucleoid segregation, physical forces are likely to play a significant role in the segregation of sister nucleoids [20, 34]. The second hypothesis postulates that some kind of stable connection between chromosomes was created at the time when the *lacZ* loci underwent DSBR-mediated physical interaction and this connection persists for several generations between the two chromosomal complexes until those DNA molecules are segregated at cytokinesis (Fig 6A). Such stability would likely require the connection to be covalent in nature and could take the form of chromosome dimerization and/or chromosomal interlinking (catenation), both of which would permit sister-*lacZ* segregation while providing a barrier to cousin chromosome segregation. The two alternate hypotheses presented here are not mutually exclusive.

## Materials and Methods

### Bacterial Strains and Growth

The genotypes of all *E. coli* strains used in this study are listed in Table S1. Cultures of *E. coli* were grown in LB growth medium at 37 °C with vigorous shaking and maintained in exponential growth phase by dilution. Cellular density was approximated by the optical density at 600 nm. Growth rates were measured using the software package fitderiv v1.03.

### Microscopy Image Acquisition

Cultures were grown in LB at 37 °C under agitation. Cells from an overnight growth were diluted into prewarmed medium to an OD_600nm_ of about 0.02 and grown for 80 minutes until an OD_600nm_ around 0.2. Then, the cultures were diluted again to an OD_600nm_ of about 0.02 and grown for 40 minutes to an OD_600nm_ lower than 0.1. A sample was taken for microscopy and mounted on a pad of 1% agarose-H _2_O for viewing under the microscope. Phase and fluorescence images were acquired at a magnification of 0.1 µm per pixel using a Zeiss Axiovert 200 widefield fluorescence microscope equipped with a Photometrics Evolve 512 EMCCD camera controlled with commercially available software (MetaMorph). For data described in Fig 1 and 4, cells were treated with 200 mg/l of chloramphenicol and 1 mg/l of DAPl prior to 2D imaging. For data described in Fig 5, 3D (z) images were acquired at 200 nm intervals. Four independent biological repeats were obtained for each sample.

### Image Processing and Analysis

For data described in Fig 5, 3D fluorescence images were deconvolved by adaptive 3D deconvolution prior to image analysis. 3D deconvolved fluorescence images were converted to 2D images by taking the maximum projection. 3D phase images were converted to 2D images by using the Best Focus function of MetaMorph. All image analyses were performed on 2D data. Cells, nucleoids, and *lacZ* foci were segmented using the cell, object, and spot segmentation algorithms of Oufti, respectively. Subpixel *lacZ* foci positions were determined by Oufti. For data in Fig 1 and 4, only cells containing either 1 or 2 nucleoids (as defined by the Oufti object segmentation algorithm) were taken into account (Fig S1C). The lengths of all segmented cells were first appended to the Oufti generated cellist structure using custom MATLAB function getExtraDataLoop, before extracting the length of segmented cells containing 1 or 2 nucleoids using custom MATLAB function getCellLengths1or2Nucleoids. Radius at mid-cell was calculated using the custom MATLAB function getMidCellWidths. Function getMidCellWidths, calculates the mid-cell width of cells meeting the following criteria: longer than the estimated average length at birth, shorter than the estimated average length at division and containing either 1 or 2 nucleoids. Mid-cell width was defined as the minimum of the central three Oufti cell width measurements. The mid-cell width of cells within 100 nm bins was averaged and data from biological repeats aggregated. Mid-cell radius was defined as half of the mid-cell width.

### Reconstructed Population Average Cell Cycles

Exponentially growing cultures were assumed to be ‘ideal’ with no cell to cell variation, with the distribution of relative cell ages given by the formula (1), [14].

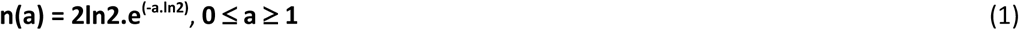

**a** is relative cell age

**n(a)** is the probability density of a cell to be of age **a**

#### Estimated average length at birth and division

Estimated average length at birth was calculated from mean length of cells with 1 or 2 nucleoids (below) using formula (2), [6]. Cells with more than 2 nucleoids were excluded in an attempt to remove filamentous cells from the sample that invalidate the assumption of an ideal age population structure. Estimated average length at division (L_d_) was defined as double the estimated average length at birth.

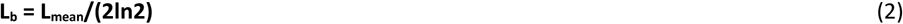

**L**_**b**_ is the estimated average cell length at birth

**L**_**mean**_ is the average cell length in the measured sample

#### Estimated time since birth

Cell length was converted to estimated time since birth using formula (3).

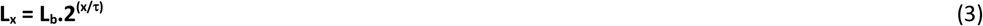

**L**_**x**_ is the cell length at time x

**L**_**b**_ is the length at birth

**τ** is the mass doubling time

#### Demographs

Reconstructed average cell cycles presented in Fig 4 were created using function avgCellCycleFromSnapshot_demograph, which was derived from the demograph function of Oufti. Briefly, cells whose length were within the estimated average length at birth and estimated average length at division and that contained either 1 or 2 chloramphenicol condensed, DAPl-stained nucleoids (above), were sampled taking account of the ideal population structure (Fig S1F) to prevent over-representation of new born cells and under-representation of cells near division. DAPl signals were summed along the long axis of each cell to give a 1-dimensional representation of the DNA density. The resultant signal for each cell was aligned along its mid-point, and each cell ranked in order of cell length. Data from the four biological repeats were aggregated.

### *oriC* Copy Number Determination by Replication Run Out

Cells grown in LB at 37 °C under agitation from an overnight growth were diluted into fresh medium to an OD_600nm_ of about 0.02 and grown for 120 minutes until an OD_600nm_ around 0.5. Then, the cultures were diluted again to an OD_600nm_ of about 0.02 and grown for 60 minutes to an OD_600nm_ lower than 0.2. Initiation of DNA replication and cell division were then blocked by the addition of rifampicin (150 µg/ml) and cephalexin (10 µg/ml), respectively. 2 ml of cells were collected, washed twice in 1 ml of PBS and resuspended in a mixture of 0.1 ml PBS and 0.9 ml ice cold ethanol. Samples were stored at 4 °C before staining DNA. To stain DNA, samples were centrifuged on a bench top centrifuge for 3 minutes before removing the supernatant. Residual ethanol was then allowed to evaporate by leaving uncapped tubes at room temperature for 20 minutes. Fixed cells were then washed twice in PBS, resuspended in DNA staining solution (10 µg/ml of propidium iodide and 100 µg/ml of RNase A), and left in the dark for 1 hour before measuring cellular fluorescence using an Apogee A50 flow cytometer. Three independent biological repeats were obtained for each sample.

In order to convert emitted fluorescence to chromosomal copy number (which indicates *oriC* copy number at the time of inhibition of replication initiation and cell division), controls are required with a known DNA content. Stationary phase cultures of cells grown in nutrient poor (M9 glycerol) media were fixed, stained and analyzed as above and the lowest (major) peak of fluorescence was assumed to correspond to 1 chromosome unit.

To determine the fraction of cells that were yet to initiate replication, two Gaussian distributions were fitted to the distributions of emitted fluorescence of object using the fitgmdist function of MATLAB after applying a threshold (150) to remove non-fluorescent particles from the analysis. For all samples, the mean value of the one of the fitted distributions was confirmed to be double the mean value of the other fitted distribution.

### Average Cell Length at Replication Initiation

To calculate the average cell length at replication initiation, the average relative cell age at replication initiation was first calculated using formula (4), [6]. This was then converted to the average cell length at replication initiation using formula (5), [6].

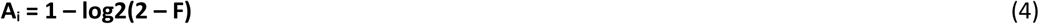

**A**_**i**_ is estimated average relative cell age at chromosomal DNA replication initiation

**F** is the fraction of cells in the sampled population that had yet to initiate chromosomal DNA replication at the time of rifampicin/cephalexin addition

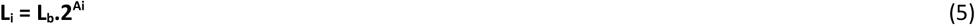

**L**_**i**_ is the estimated average cell length at chromosomal DNA replication initiation

**L**_**b**_ is the estimated average cell length at birth

**A**_**i**_ is estimated average relative cell age at chromosomal DNA replication initiation

### Analysis of Chromosomal DNA Replication

#### Chromosomal Marker Frequency Analysis

Cells from an overnight growth were diluted into fresh LB medium to an OD_600nm_ of about 0.02 and grown at 37 °C under agitation until an OD_600nm_ of about 0.2, after which the cultures were diluted again to an OD_600nm_ of about 0.02 and grown to an OD_600nm_ between 0.25 and 0.3. Chromosomal DNA was extracted from cultures of cells at both mid-exponential growth phase and (non-replicating) stationary growth phase using the Wizard genomic DNA extraction kit (Promega), according to manufacturer’s instructions. An additional RNase step was performed using RiboShredder RNase Blend (Epicentre Biotechnologies). Three independent biological repeats were obtained for each sample. Library preparation and Illumina Solexa sequencing were performed on these DNAs by Edinburgh Genomics (University of Edinburgh) who subsequently provided paired-end reads with adapter sequences removed. Reads were aligned to the DL1777 draft genome sequence (Table S2) using the BurrowsWheels Alignment software BWA-MEM and the number of reads mapped to each bp of the genome was quantified using SAMtools (mpileup). To account for differences in read depth between samples, the number of mapped reads for each base on the reference genome was normalized by the total number of mapped reads for that sample. Values for exponential growth samples were then normalized against values obtained for stationary phase growth values to account for potential sequencing biases. This normalization assumes an equal copy number of each locus across the chromosome in stationary phase (‘non replicating’) cultures. Normalized read values were converted to estimated average cellular copy number using the average cellular *oriC* copy number calculated by replication run out experiments (above).

#### Replication Rates and Position of Termination

Replication rates were derived from Linear regression on log2 transformed normalized MFA data (above) using formulae (6) and (7), [17]. For these calculations, the left replichore was defined as genome coordinates 1,580,001 to 3,835,000 and the right replichore was defined as genome coordinates 4,324,500 to 1,277,500. The coordinate of replication termination was defined as the point of intercept between the lines of best fit to the left and right replichores.

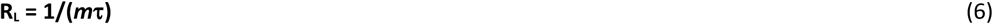

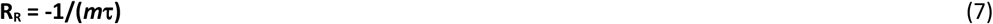

**R**_**L**_ is the replication rate of the left replichore in bp/s

**R**_**R**_ is the replication rate of the right replichore in bp/s

***m*** is the gradient of the line of best fit to the log2 transformed normalized MFA data

**τ** is the mass doubling time in seconds

#### C-time

To calculate the time to complete chromosomal DNA replication, the replication rate for each replichore (above) was multiplied by the number of bp replicated (the distance between *oriC* mid-point and the co-ordinate of replication termination, above).

#### Average Cell Length at lacZ *Replication*

The time required for a replication fork to reach *lacZ* (~15 min) was calculated using the distance of *lacZ* from *oriC* (1,081,626 bp) and the replication rate of the right replichore (above) and added to the estimated average time between birth and replication initiation (derived from the relative age at replication initiation (A_i_) and the generation time). Since *lacZ* was found to replicate in the subsequent generation, one generation time was subtracted from this value to give an estimated time since birth (in the subsequent generation) at *lacZ* replication. The average time between birth and *lacZ* replication was converted to average cell length at *lacZ* replication using formula (3).

## Acknowledgements

We are grateful to Nancy Kleckner for helpful discussions and financial support to M.A.W, and to Edinburgh Genomics for the whole genome sequencing. Edinburgh Genomics has been partly supported through core grants from Natural Environmental Research Council [R8/H10/56]; Medical Research Council [MR/K001744/1]; and Biotechnology and Biological Sciences Research Council [BB/J004243/1].

## Author Contributions

Conceptualization, M.A.W. and D.R.F.L.; Methodology. M.A.W. and D.R.F.L.; Software, M.A.W.; Formal Analysis, M.A.W; Investigation, M.A.W., E.D., and M.A.L.; Resources, D.R.F.L.; Writing - Original Draft, M.A.W. and D.R.F.L.; Writing - Review & Editing, M.A.W., E.D., M.A.L. and D.R.F.L.; Visualization, M.A.W. and D.R.F.L.; Supervision, D.R.F.L; Project Administration, D.R.F.L; Funding Acquisition, D.R.F.L.

## Funding

Medical Research council [G0901622 and MR/M019160/1 to D.R.F.L]; Human Frontiers Science Program [LT000927/2013 to M.A.W.]. National Institute of Health [GM044794 awarded to Nancy Kleckner].

## Figure Legends

**Supplemental Figure 1. Related to Figure 1. DSBR causes a SfiA-independent increase in average cell length at birth. A)** Asynchronous cultures of *E. coli* were grown in rich growth media at 37 °C and maintained in exponential growth phase by serial dilution. Population growth was measured as the optical density at 600nm (OD_600_). **B**) Average mass doubling time derived from rate of change of OD_600_. **C)** Number of cells analyzed following chloramphenicol-DAPl treatment. **D)** Mean cell length of cells containing either 1 or 2 chloramphenicol condensed DAPl stained nucleoids. For **A, B** and **D**, error bars show standard error of the mean, n = 4 independent cultures. **E)** Age structure of ideal asynchronous population with twice as many newborn cells as cells on the cusp of division. **F)** Expected distribution of relative cell lengths in an ideal asynchronous population assuming exponential growth. For **E** and **F**, data were normalized to approximate probability distribution functions (PDF). **G)** Histograms of measured cell lengths in asynchronous cultures with annotated estimated average length at birth (L_b_) and estimated average length at division (L_d_) calculated using the measured mean length of cells with either 1 or 2 chloramphenicol condensed nucleoids and assuming an ideal asynchronous population of cells. The data from four independent experiments for each *E. coli* strain were aggregated. **H)** Cell radius at mid-cell as a function of estimated time since birth, with estimated time since birth calculated as a conversion of measured cell length using the measured average mass doubling rate and estimated length at birth. For both *sfiA*^+^ and Δ*sfiA* strains, the data from the three control strain backgrounds not undergoing DSBR (SbcCD^+^ *lacZ*^+^, SbcCD^-^ *lacZ*^+^ and SbcCD^-^ *lacZ*::palindrome) were averaged for clarity in the plots. No difference was detected between the three control strains in either *sfiA*^+^ or Δ*sfiA* backgrounds.

**Supplemental Figure 2. Related to Figure 3. DSBR alters the chromosomal DNA replication profile without affecting the time required to complete DNA synthesis. A**) The distribution of mapped sequencing reads across the genome for each of the biological repeats of the four strains. **B**) The mean MFA for three independent cultures of strain DL2859 (*sbcCD*^+^ *lacZ*::palindrome) experiencing DSBR at *lacZ* normalized against the mean MFA for three independent cultures of each of the three control strains (1777: *sbcCD*^+^ *lacZ*^+^; 2151: *sbcCD*^-^ *lacZ*^+^; 2874: *sbcCD*^-^ *lacZ*::palindrome). **C**) Linear regression of MFA results used to calculate the replication rates of the left and right replichores shown in Fig 3C. **D**) Intercept of the lines of best fit to the left and right replichores, used to calculate the predicted location of replication termination (Fig 3D). A moving mean of 10,000 bp was applied to the read counts to create the plots shown in **A** - **C**.

**Supplemental Figure 3. Related to Figure 5. The locus undergoing DSBR dwells at future division planes.** Spatial distribution of *lacZ* adjacent LacI-CFP and TetR-YFP foci along the long axis of cells undergoing DSBR at *lacZ* (SbcCD^+^ Palindrome^+^, red), or not.

**Supplemental Table 1. Strain list**. A list of *E. coli* strains used in this study.

**Supplemental Table 2. Deposited Data**. A list of publicly accessible data generated in this study.

**Supplemental Table 3. Software list**. A list of software/functions used and generated in this study.

